# Non-invasive Biomechanical Characterization of Embryos Using Microfluidic Cantilevers

**DOI:** 10.64898/2025.12.02.691595

**Authors:** Irene C. Turnbull, Tai De Li, Pedro Sanabria, Aimee Stablewski, Angelo Gaitas

## Abstract

Embryonic development is intricately regulated by mechanical properties such as stiffness, which influence developmental viability and implantation success – factors critical in assisted reproductive technologies (ART). Traditional embryo evaluation relies predominantly on morphology, lacking quantitative mechanical parameters that could enhance selection accuracy. Recent studies indicate that the stiffness (elasticity) of the zona pellucida (ZP) – the glycoprotein-rich extracellular matrix surrounding mammalian oocytes and embryos – correlates with embryo quality and developmental potential. However, current biomechanical characterization techniques – including micropipette aspiration, atomic force microscopy (AFM), microtactile sensors, and MEMS-based systems – either pose risks of mechanical damage or involve complex, time-consuming procedures unsuitable for clinical settings. Here, we introduce a novel approach leveraging fluidic force microscopy cantilevers to non-invasively evaluate embryo biomechanics. Our proof-of-concept study demonstrates rapid, precise stiffness profiling of intact mouse embryos (specifically ZP elasticity) in under one minute per embryo. Using gentle microsuction attachment with no chemical adhesives or rigid immobilization, the method preserves embryo integrity while providing reproducible elasticity measurements. This method combines the precision of AFM with minimal invasiveness, offering a promising new quantitative biomechanical indicator to augment clinical embryo assessment and paving the way for broader applications in reproductive biology.

## Introduction

Embryonic development is governed by intricate mechanical cues – including tension, compression, and elasticity – that orchestrate cell positioning, lineage decisions, and tissue morphogenesis even before overt genetic patterning ensues (Maître et al., 2016; Eisenstein, 2024). Among these, elastic mechanical properties play a critical role in embryo development and implantation potential, which is especially relevant for assisted reproductive technologies (ART)^1,2^. The animal industry has benefited extensively from ART, including in vitro fertilization (IVF), aiding in the efficient reproductive performance in livestock and improvement of genetic quality^3^; these tools are also applied in basic research, the generation of genetically modified animals and the preservation of endangered species ^4,5^. But IVF success rate is hampered by embryo quality challenges^6^. The zona pellucida (ZP), a glycoprotein-rich extracellular coat encasing mammalian oocytes and early embryos, is crucial for fertilization (facilitating sperm binding and penetration), prevention of polyspermy via post-fertilization “zona hardening”, and protection during preimplantation development^7,8^. Indeed, any inability of sperm to penetrate the ZP can cause fertilization failure^7^. Current embryo selection in IVF is largely based on subjective morphological grading under a microscope ^9,10^. While useful, morphology alone does not capture biophysical properties – such as the stiffness of the embryo’s encapsulating ZP – that are fundamental to developmental competence ^11,12^; this has led to a growing emphasis on identifying additional markers that can improve ART success rates^13^. Recent evidence suggests that ZP biomechanical properties could serve as objective markers of oocyte and embryo quality. For example, measuring elasticity of the ZP using a microtactile sensor showed distinct mechanical differences between immature and mature oocytes, and this can aid in the selection of high quality oocytes for ART ^14,15^. Notably, the thickness of the ZP will impact the success rate of IVF, the implantation process, and pregnancy rates ^6^. These findings support the idea that ZP elasticity is a meaningful indicator of oocyte/embryo viability that could augment existing selection criteria^16^.

Several methods have been explored to quantify embryo mechanical properties, each with advantages and limitations. Micropipette aspiration can directly measure viscoelastic responses by suction of the embryo/ZP, and has been shown to differentiate mechanical hardness before and after fertilization ^8^. For instance, one study using micropipette aspiration found that fertilization causes a significant increase in ZP stiffness (consistent with the zona hardening cortical reaction)^8^. Another study, also applying a micropipette aspiration system demonstrated its usefulness in evaluating embryo stiffness, where the aspiration depth of the embryo into the pipette under a prescribed pressure is fit into a mechanical model; these data were predictive of embryo viability proving its usefulness in embryo selection for ART ^17^. However, some micropipette methods are relatively invasive and labor-intensive: they involve physically deforming the embryo with a pipette, which risks mechanical stress or even microscale damage to the ZP or cells, may affect levels of gene expression, and they require careful modeling to extract a Young’s modulus ^8,17,18^. Atomic force microscopy (AFM) has been applied to measure oocyte/embryo stiffness at high spatial resolution ^19,20^. AFM indentations can reveal layered mechanical responses of the ZP (e.g. distinguishing a stiff outer layer vs. softer inner layer) and have confirmed trends such as ZP softening with oocyte maturation ^21^. AFM indentation measurements showed that the mechanical properties of oocytes are related to their quality and ability to develop into viable blastocysts after fertilization; ZP stiffening in aged oocytes (6-hour vs 1-hour post thaw), along with decrease in viscous energy, correlated with lower IVF yields ^19^. Yet, AFM typically requires the sample to be immobilized on a substrate (often via adhesion) to avoid slippage during probing. This can introduce chemical adhesives or abnormal constraints that may alter the embryo’s environment. Moreover, standard AFM probes and controllers are not designed for the curved, free-floating embryo, making it challenging to obtain reproducible results without risking perturbation or even penetration of the embryo. Microtactile sensors and MEMS-based devices have also been developed to assess ZP mechanics. For example, Murayama *et al.*^15^ used a microfabricated tactile sensor to measure mouse ZP hardness, finding that a fertilized mouse embryo’s ZP was about 2.7-fold stiffer than an unfertilized oocyte’s ZP ^8^. While such devices can handle whole embryos and provide quantitative stiffness data in the kPa range, they often necessitate complex setups or significant embryo deformation, and are not yet practical in IVF workflow ^15^.

To address these challenges, we apply Fluidic AFM (FluidFM) – a technique that integrates microfluidics into an AFM cantilever – as a gentle, non-invasive platform for whole-embryo biomechanical characterization. FluidFM was introduced by Meister *et al.* (2009)^22^ as an AFM with microchanneled cantilevers, allowing precise fluid control through the cantilever ^22^. In this approach, a tiny aperture at the cantilever tip enables suction or dispensing at localized spots. For our purposes, the hollow cantilever can act as a microscopic “vacuum hand” to gently grasp and release embryos by suction, eliminating the need for adhesives or rigid clamping of the embryo. This gentle approach minimizes mechanical stress and preserves sample viability. FluidFM has already proven effective in various delicate bio-applications, including single-cell isolation and adhesion measurements, injection of biomolecules into cells, and even weighing single cells in liquid ^23–27^. These successes underscore its suitability for handling fragile biological specimens. Despite extensive use of FluidFM in cell biology, its utility for whole embryo mechanics remains unexplored.

In this proof-of-concept study, we demonstrate that FluidFM can rapidly measure the elastic modulus of intact mouse embryos’ ZP in a non-invasive manner. We present the first quantitative stiffness measurements of mouse embryos obtained via FluidFM, and compare the results to prior AFM-based studies of ZP mechanics. We further discuss how this method could enhance embryo selection in IVF by providing a new biomechanical metric, and outline future integration of simultaneous mass measurements to create a more comprehensive embryo evaluation tool.

## 3. Materials and Methods

### Embryo Preparation

Cryopreserved mouse embryos (late preimplantation stage, wild-type C57BL/6 strain) were obtained on dry ice and stored in liquid nitrogen. For each experiment, embryos were thawed using a standard rapid thaw protocol (modified from Renard & Babinet 1984^28^). In brief, straws containing vitrified embryos were retrieved from liquid nitrogen and held in room air for ∼40 seconds before being plunged into a +25 °C water bath until ice had just melted (∼5–10 seconds). The contents of each straw were expelled into culture medium (M2 media, Millipore Sigma) and the embryos were allowed to rehydrate for 5 minutes, during which they visibly shrank and then re-expanded to normal morphology. Embryos were then washed through a series of 10 drops of fresh M2 medium to remove cryoprotectant. All steps were conducted at room temperature under sterile conditions. The recovered embryos were visually inspected to confirm intact morphology and then maintained in M2 medium at ambient temperature during mechanical testing. All procedures were carried out in compliance with relevant ethical guidelines for animal research.

### FluidFM-AFM Instrumentation and Mechanical Testing Procedure

We utilized a Flex-Bio AFM system (Nanosurf AG, Switzerland) combined with a FluidFM microchanneled cantilever (Cytosurge AG, Switzerland) mounted on an inverted optical microscope (Zeiss Axio Observer) for alignment and observation. The FluidFM probe consists of a silicon nitride microcantilever with an integrated microfluidic channel and a circular aperture at the end of the cantilever (aperture diameter ∼8 µm). This setup was enclosed in an environmental chamber to minimize evaporation and maintain stable conditions. Prior to measurements, the cantilever’s spring constant was calibrated with the Sader method ^29^. The fluidic system was connected to a pressure controller/pump (Cytosurge) capable of applying pressures from −800 to +1000 mbar to the cantilever’s interior channel. For our embryo tests, we used a gentle negative pressure (approximately –100 mbar) to pick up and hold each embryo via suction at the cantilever aperture, and a positive pressure to quickly release the embryo after testing. The entire procedure was performed with the embryos submerged in M2 medium at +37 °C, in a Petri dish.

The FluidFM-based measurement involved the following steps (illustrated in **figure 1**):

1. **Approach & Capture:** The cantilever was maneuvered until the aperture made contact with the embryo’s ZP. A gentle suction (negative pressure) was applied, causing the embryo to adhere to the aperture (without penetrating the ZP).
2. **Compression:** With the embryo attached, the cantilever was driven downward toward the substrate (dish) in a force spectroscopy mode. The embryo, being held by the cantilever, was thus sandwiched between the cantilever (on top) and the dish (on bottom). The AFM recorded force–displacement curves during this compression. We limited the maximum applied force to avoid over-compressing the embryo, ensuring deformations remained small (<5% of embryo diameter) to stay in the elastic regime.
3. **Measurement:** From the force–displacement curve, the effective stiffness of the embryo/ZP was determined. We modeled the system as an elastic sphere (the embryo) being compressed between two rigid surfaces (the cantilever and the dish). A modified Hertz contact model was used to extract the Young’s modulus (E) of the embryo’s ZP from the force–displacement data (details below).
4. **Release & Repeat:** After obtaining a force curve, the cantilever was raised and a slight positive pressure was applied to cleanly detach (release) the embryo from the cantilever tip. The same cantilever was then reused to capture the next embryo in sequence, repeating steps 1–3 for each of the embryos.

**Figure 1.**
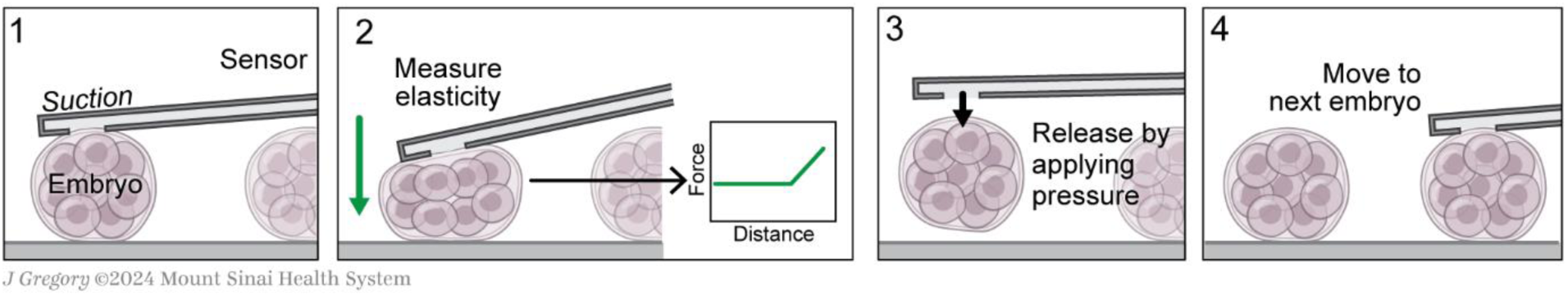
Schematic of the Fluid-FM based measuring process. 1) The cantilever contacts the embryo while we apply suction. 2) The embryo is compressed while the AFM records the force-distance curve and the elasticity of the embryo is measured. 3) Applying positive pressure to release the embryo. 4) The same cantilever is moved to the next embryo to repeat the process.

### Duration of measurements

The process of identifying and attaching the embryo takes <1min, the approach to the bottom of the dish takes ∼30 seconds. Each force-distance curve measurement is completed in ≤2 seconds. The cantilever is moved at a distance of 10µm at a ramp rate of 10µm s⁻¹ (1 second approach and 1 second retract). The post-contact loading segment was displaced by ∼2 µm in distance and lasted ∼0.2 seconds. We acquired three curves per embryo, separated by 20-30 seconds, then we retracted the probe (∼10-30 seconds) and we applied positive pressure to release the embryo. After completing each measurement and before release, we kept the embryo under observation attached to the probe for several minutes to screen for delayed damaging effects due to the measurement.

### Elastic Modulus Calculation

We employed a Hertz-model-based analysis to calculate the apparent Young’s modulus of the embryo’s ZP from the recorded force–displacement data (**Fig. 2**). Because the embryo was in contact with two surfaces (cantilever tip and dish), we assumed, as a first approximation, that the total displacement Δd was split equally between the top and bottom contacts (i.e. each side displaces the ZP by δ ∼ Δd/2). The force F_N top_ measured by the cantilever is the same force compressing the embryo against the dish F_N bottom_ (**Fig. 2**). For a spherical elastic object of radius *R* compressed by a rigid flat surface (Hertz model for sphere-flat contact), the force and displacement are related by:

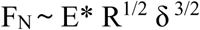

where E* is the effective elastic modulus (for our purposes, essentially the ZP’s modulus, since the embryo’s interior is much softer) and δ is the displacement at that contact. In our dual-contact scenario, substituting δ = Δd/2 yields

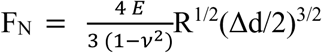

We treated the embryo as a homogeneous sphere and assumed a Poisson’s ratio ν approx. 0.5, as is typical for soft biomaterials under compression ^8^. The radius R of each embryo was measured individually from microscope images (∼50–60 µm radius, consistent with mouse embryo ZP diameter) as described in ^26^. Using the above relation, we derived the *E* for that embryo from the force-distance graphs using AtomicJ^30^. For details of the fitting procedure and detailed derivation, refer to **Supplementary File S1 and Fig. S1.**

**Figure 2.**
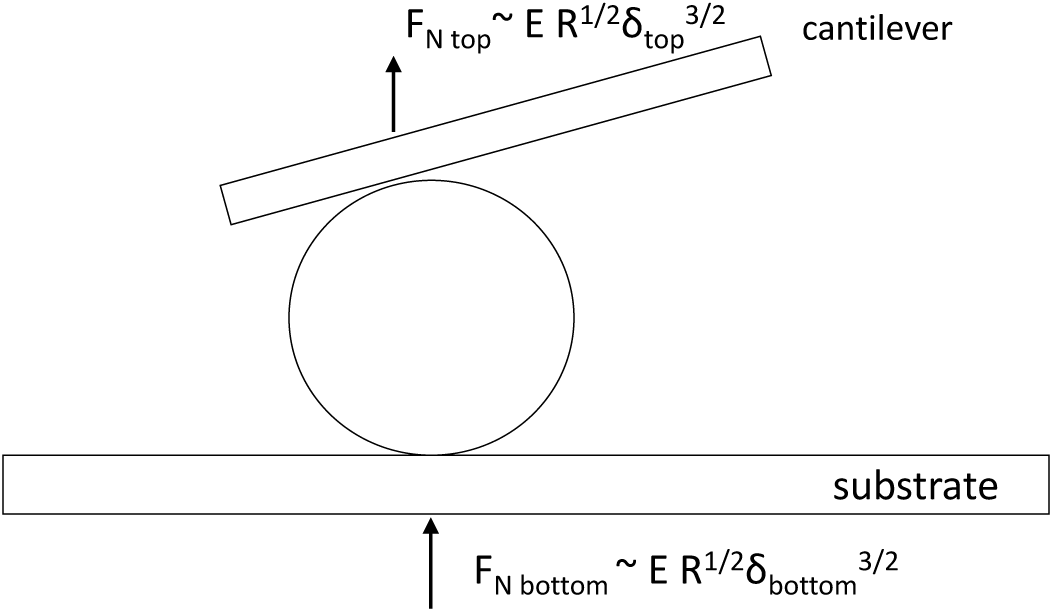
The mechanical model used to extract the apparent Young’s modulus of the embryo’s zona pellucida (ZP) from the force–displacement curves. The embryo (sphere, radius *R*) is compressed between two rigid surfaces: the cantilever (top plate) and the substrate dish (bottom plate). Solid vertical arrows indicate the normal forces applied at each interface (F_N top_ and F_N bottom_). Because the compression is symmetric, each interface contributes one-half of the total displacement (δ ≈ Δd / 2).

## Results

Using the FluidFM approach, we successfully measured the ZP elasticity of individual mouse embryos without any visible damage or disruption to the embryos. Each measurement (from initial capture to release) was completed in under 1 minute, demonstrating the rapid throughput potential of this method. In all cases, the embryos remained intact and morphologically normal after release, confirming the non-invasiveness of the technique.

Each embryo was measured multiple times (3 force curves per embryo) to assess reproducibility. Throughout testing, the embryos remained viable-looking and freely floating upon release, indicating minimal physical perturbation by the measurement. A representative force–distance curve for a whole embryo is shown in **figure 3a**. A still image at the time of capture and after release is shown in **figures 3b and 3c**, respectively. For additional time-course panel of another embryo refer to **Fig. S2** and force–distance indentation curves and Hertz fits refer to **Fig. S3** and **Table S1.** The image of the embryo before and after release for comparison show that the embryo’s morphological appearance remains intact after the measurement procedure. **Video 1** shows the release.

**Figure 3.**
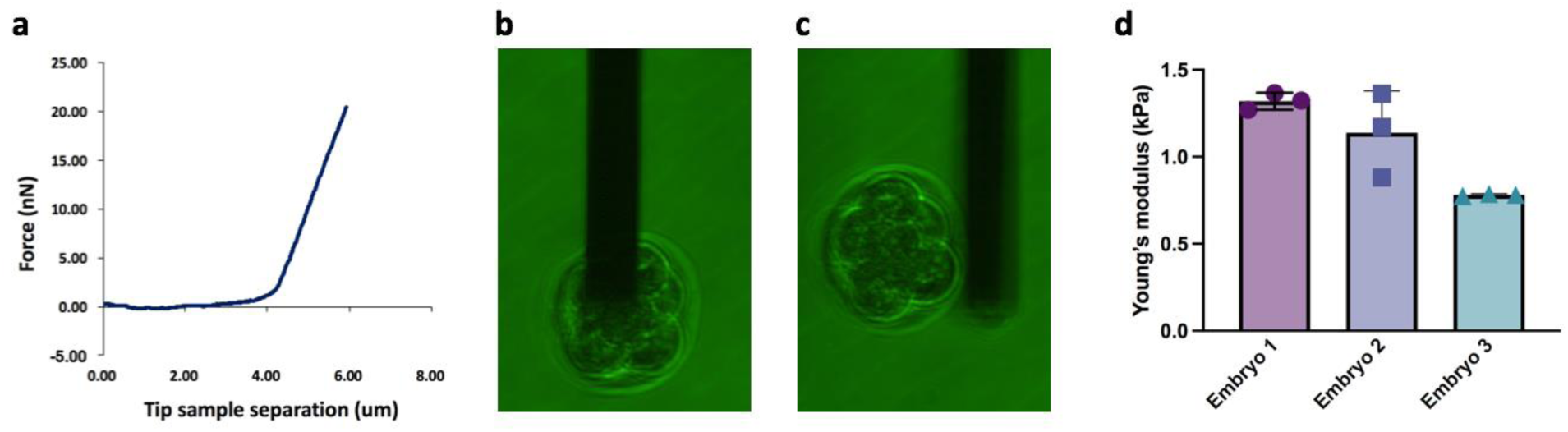
Data acquisition and results. a) Representative graph of a whole embryo force curve. (b) Picture of embryo being attached using suction and (c) then being released with positive pressure. d) Summary of Young’s modulus measurements for three individual mouse embryos using FluidFM-AFM. Each bar represents the mean apparent Young’s modulus (E) calculated from N = 3 indentations per embryo using a modified Hertz model. Error bars denote the standard deviation of the measurements. Embryo 1: mean E = 1319 ± 48 Pa, Embryo 2: mean E = 1139 ± 241 Pa, and Embryo 3: mean E = 781 ± 6 Pa.

The stiffness results for the three embryos are summarized in **figure 3d**. All three embryos exhibited ZP Young’s modulus values on the order of 10^2^ Pa. All measurements are of the same order of magnitude, suggesting good reproducibility of the method and a relatively consistent ZP stiffness for these embryos. The small differences observed between embryos (Embryo 3 being slightly softer than Embryos 1 and 2) could be due to biological variability in ZP composition or cryopreservation effects; with our small sample size, we cannot draw firm conclusions, but this hints that the technique may be sensitive enough to detect real stiffness differences between embryos. Across all embryos the calculated Young’s moduli fell in a range of ∼0.78 to 1.37kPa.

## Discussion

### Comparison with AFM-Based Measurements

The elasticity values obtained with FluidFM-AFM are in good agreement with the order of magnitude reported by previous AFM studies of mammalian oocytes. AFM indentation on human oocytes found that the outermost layer of the ZP has an elastic modulus on the order of ∼100 Pa, whereas deeper ZP regions and whole-oocyte responses were a higher (several hundred Pascals)^21^. In contrast, micropipette aspiration techniques, which deform the entire ZP, tend to report higher apparent stiffness – usually in the single-digit kPa range ^31^. This discrepancy has been noted in the literature and is attributed to differences in contact area and deformation: AFM probes a small local patch of the outer ZP, which may be more compliant, whereas micropipette aspiration engages a large portion of the ZP and underlying cell, yielding a higher effective modulus ^21^. Our FluidFM method lies between these two extremes: by compressing the whole embryo gently between two surfaces, we are probing a broader area of the ZP than a point AFM tip, but with smaller strain than a full micropipette suction. Consequently, our measured modulus (∼∼0.78 to 1.37 kPa) is intermediate between AFM nano-indentation (∼0.1 kPa for outer ZP) and micropipette results (1–10 kPa)^21^ and (3.1 ± 0.4 kPa)^31^. This is a reassuring consistency, suggesting that FluidFM provides a physically meaningful measure of ZP stiffness.

Furthermore, our proof-of-concept results hint at the ability to discern mechanical differences between embryos. Although all three embryos were nominally similar, Embryo 3’s ZP was ∼30-40% softer. Such differences might correlate with embryo quality or developmental stage. Prior AFM studies support this idea: Papi *et al.* (2010)^7^ showed that in bovine oocytes, the ZP’s mechanical behavior shifts from purely elastic in immature eggs to more compliant (and partially plastic) in mature eggs, then stiffens again after fertilization due to zona hardening ^7^. In our measurements of post-fertilization embryos, all had undergone the cortical reaction (as they developed to multi-cell stage), so a relatively elevated stiffness is expected. Murayama’s microtactile sensor measurements in mice indeed quantified that a fertilized embryo’s ZP (∼22.3 kPa)^32^ was substantially stiffer than an unfertilized oocyte’s ZP (∼8.3 kPa)^8^. While the absolute values differ (their sensor likely measured a different regime of deformation), the qualitative trend holds: successful fertilization induces ZP stiffening. Our method could potentially be used to confirm if an embryo has properly undergone zona hardening (an indicator of normal fertilization), or to detect aberrations (too soft could indicate failed or abnormal cortical granule release ^21^.

### Implications for IVF Embryo Selection

The ability to rapidly and non-invasively gauge embryo stiffness could add a valuable dimension to embryo assessment in ART. Currently, embryo selection relies on morphology and sometimes genetic screening, but these do not capture the biophysical state of the embryo. A mechanical readout like ZP elasticity might reflect underlying developmental competence – for instance, an overly soft ZP might signal an embryo that had an incomplete zona reaction or a developmental anomaly, whereas an excessively stiff ZP might impede hatching. Some studies have suggested that embryos which are too rigid or too soft are less likely to be viable and have lower implantation potential^16,17^. Thus, an optimal range of ZP stiffness might exist, and measuring this parameter could refine the selection of embryos with highest likelihood of success. Unlike subjective morphology scoring, stiffness is a quantitative metric. Moreover, it may provide early indications of viability even when morphology looks identical. For example, two blastocysts of similar appearance might have different ZP mechanics due to differences in their biochemical composition; our technique could discern that. Over time, accumulating data on embryo stiffness versus clinical outcomes could establish thresholds or patterns (perhaps in combination with other factors like embryo metabolism / growth rates) to guide clinicians. It is noteworthy that our method is inherently gentle – we observed no detriment to the embryos from brief FluidFM handling. The use of a mild suction to hold the embryo is analogous to procedures already routine in IVF labs (e.g. holding an oocyte with a suction pipette during intracytoplasmic sperm injection), which have proven safe. Indeed, we and others have shown that applying suction through a microchanneled cantilever does not harm cells nor alter the mechanical reading ^26,27,33^. This suggests that integrating such a measurement step in an IVF laboratory could be done without compromising the embryo.

### Advantages, Future Outlook and Limitations

Our approach offers several key advantages over traditional mechanical testing of embryos. There is no need to remove or damage the ZP, attach beads, or subject the embryo to large deformations. The entire process is gentle and quick, which is crucial for preserving embryo viability. Unlike some AFM methods that require adhering the embryo to a substrate, the FluidFM uses suction to hold the embryo only transiently, leaving no residues. We performed <1 min per embryo measurements; it is expected that with automation, a series of FluidFM cantilevers could potentially measure dozens of embryos within an hour, which is compatible with clinical IVF workflows that handle many embryos. The same cantilever can pick and measure multiple embryos sequentially, reducing cost per test and time. In addition to stiffness, the FluidFM could be used to measure other physical parameters. Three-dimensional live imaging performed on preimplantation embryos is an attractive non-invasive methodology to evaluate embryo changes, however, the measured parameters were not predictive of successful embryo transfer ^34^. Overall, this technique bridges a gap between high-precision AFM and practical embryo handling tools, offering a new way to obtain biomechanical insights in a clinically relevant manner.

Beyond stiffness, an embryo’s mass is another critical biophysical parameter. During in-vitro embryo production (IVP), variables such as maternal diet and the quality of frozen–thawed sperm can affect fertilization efficiency and subsequent development ^35,36^. Optimizing these factors has been shown to yield embryos with a larger inner cell mass—a feature strongly associated with higher implantation and pregnancy rates^36^. The ability to measure embryo mass non-invasively could provide insight into its growth rate. Traditional methods to weigh single cells or embryos (like resonant mass sensors) have limitations as discussed. Sone et al. demonstrated a holder-type piezoresistive cantilever for positioning single mouse embryos and estimating mass via resonance-frequency shifts in liquid^37^. However, the lack of rigid fixation lets buoyancy and hydrodynamic coupling dominate—causing systematic underestimation—while manual placement is low-throughput and potentially stressful. These drawbacks are mitigated by fluidic AFM, which provides gentle, secure capture/positioning in liquid and enables more stable, scalable measurements. Therefore, our platform can be extended to perform mass measurements by exploiting resonant frequency shifts of the cantilever with the embryo attached. Our group has recently demonstrated that microchanneled AFM cantilevers can act as extremely sensitive weigh scales for single cells in media, achieving mass resolution on the order of 10 picograms ^26,27^. This is sufficient to detect ∼0.01% changes in embryo’s mass, which could track subtle growth changes. To apply this to embryos, a specialized microfluidic cantilever can be designed that allows us first to measure its resonant frequency and then proceed to perform the indentation for elasticity. Such a dual-modality device would provide a more comprehensive biomechanical profile. We anticipate that combining these metrics could further improve embryo selection.

This study is a first proof-of-concept with a small sample size, so we could not correlate stiffness with developmental outcomes. In future studies, larger cohorts of embryos will be needed to statistically validate stiffness as a predictive marker. We also focused on one developmental stage; the technique should be evaluated at other stages (e.g. oocyte, zygote, blastocyst) to map how mechanics evolve through development. Although we detected no obvious harm, the next steps include: a) culturing the tested embryos to the blastocyst stage and carry out detailed genetic analyses - to verify definitively that the measurement procedure has no adverse effect on viability; and b) to perform animal transfers soon after measurement to compare stiffness with developmental outcomes.

## Conclusions

We have presented a novel application of fluidic AFM cantilever technology to characterize the biomechanical properties of intact embryos in a rapid and non-invasive manner. This proof-of-concept study measured the elastic modulus of the zona pellucida in mouse embryos using gentle microfluidic suction for handling. The results showed that FluidFM can reproducibly quantify embryo stiffness within minutes per sample, without causing damage. The measured ZP stiffness values were consistent with previously reported ranges and reflected known biological trends, underscoring the validity of the approach. By providing a quantitative mechanical readout, this technique could augment current embryo evaluation methods and improve the selection of high-quality embryos for transfer in ART. Looking ahead, the integration of mass measurement capabilities and further validation in clinically relevant models will pave the way for FluidFM-based biomechanical profiling to become a valuable tool in reproductive medicine. Ultimately, non-invasive biomechanics could help bridge the gap between traditional morphology-based selection and the underlying developmental potential of embryos, contributing to higher success rates in assisted reproduction.

## Acknowledgments

The authors gratefully acknowledge Dr. Christos Coutifaris (University of Pennsylvania, Philadelphia, PA, USA) for his insightful discussions regarding embryo elasticity measurements, which helped shape the experimental design and interpretation of our results. The authors gratefully acknowledge Jill K. Gregory, MFA, CMI, Associate Director of Instructional Technology, Icahn School of Medicine at Mount Sinai for illustrations in Figure 1. This work was supported by the National Science Foundation under award number 2344530, the content is solely the responsibility of the authors and does not represent the official views of the National Science Foundation. The Bio-AFM supported by NIH 1S10OD030401-01A1 in the Surface Science Facility of ASRC – CUNY has been utilized for optimizing the experiment parameters.

## Supplementary material

**Video 1.** Demonstration of embryo being released with positive pressure.

## Supplementary Information

### Supplementary File S1. Fitting procedure

Contact points were identified automatically in AtomicJ using the model-based estimator. We imported the data files from the Nanosurf AFM into AtomicJ for analysis. We analyzed the approach of each curve with the Paraboloid (Hertz) model (non-adhesive, linear elastic) and Poisson’s ratio ν = 0.5. For contact detection we used the Robust focused-grid search “Based on contact model,” and fits were performed with Robust (HLTS) regression (coverage ∼85–90%). This procedure determines the contact point as the position that yields the best agreement with the Hertz model while keeping the pre-contact baseline flat; the resulting contact is then used to compute indentation and fit the modulus. Because the embryo is compressed between two rigid surfaces, the total post-contact approach Δd is shared by the two contacts. Our derived dual-contact relation is (equation S1)

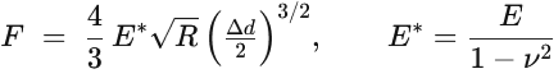

AtomicJ’s Paraboloid (Hertz) form uses the standard single-contact expression (equation S2):

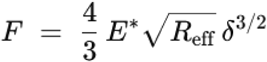

where δ is the indentation variable supplied to the model.

To use full post-contact displacement as AtomicJ’s indentation δ=Δd and still match our dual-contact physics, we set an effective radius (equation S3):

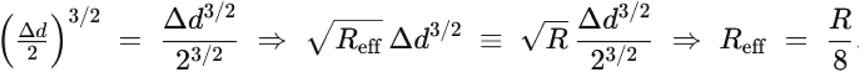

Accordingly, in AtomicJ we implemented the dual-contact geometry by specifying an effective spherical radius R_eff_=R/8 while fitting the full post-contact displacement. The embryo radii were: Embryo 1: R=54.5 μm, Embryo 2: R=49.78 μm, Embryo 3: R=50.5 μm, yielding R_eff_ values of 6.81 µm, 6.22 µm, and 6.31 µm, respectively. Representative fits for each measurement are shown above.

---

**Figure S1.**
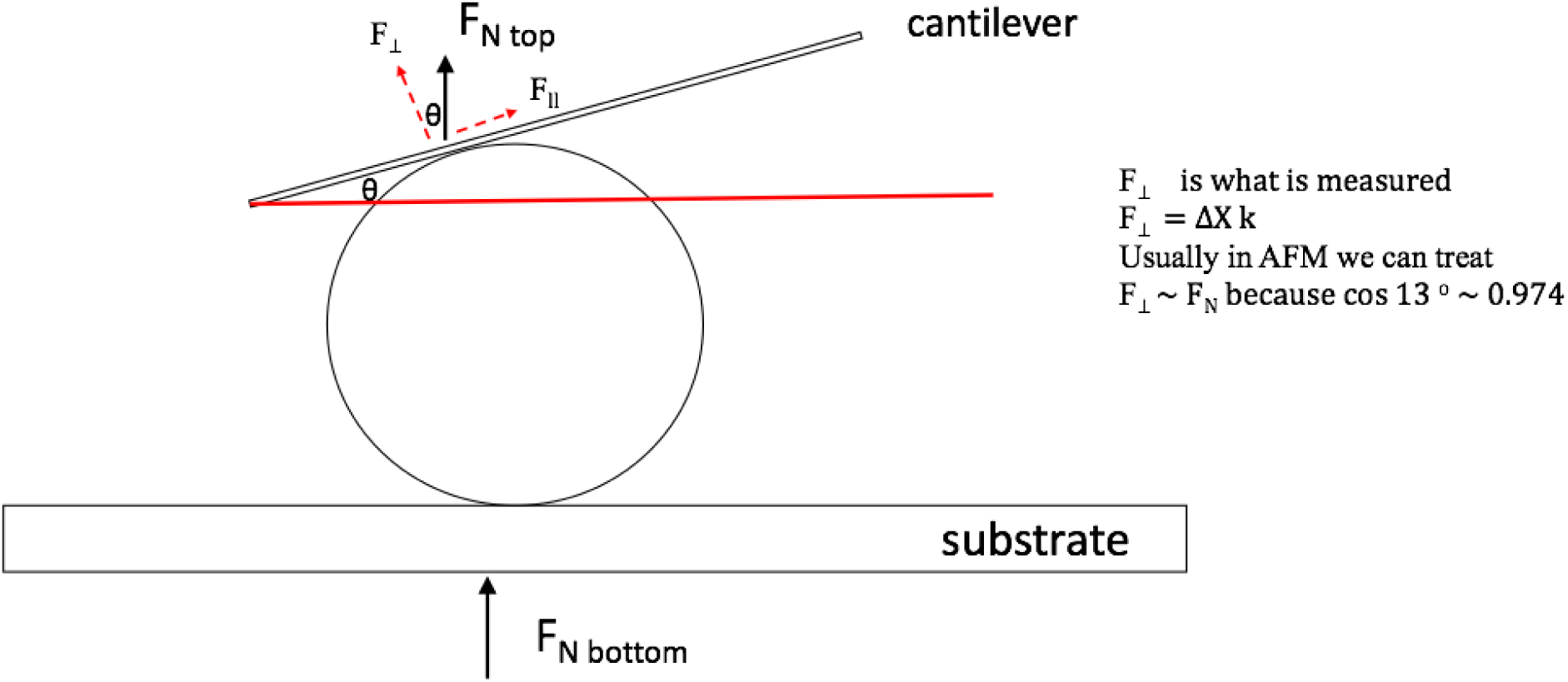
Schematic of the derivation. In our analysis, the measured cantilever load can be decomposed into components normal and parallel to the embryo surface. The normal component is the one that enters the Hertz model. F⊥=F_top_ cos θ (equation S4). For geometry θ≈13^∘^, hence Cos θ ≈ 0.974. The resulting correction is therefore 2.6%. Because this is small relative to other experimental uncertainties (e.g., baseline drift, CP placement), our primary analysis treats F_⊥_≈F_top_. While planar probes can eliminate tilt by construction (as in L. Andolfi, S.L.M. Greco, D. Tierno, R. Chignola, M. Martinelli, E. Giolo, S. Luppi, et al., Acta Biomater. 94, 505-513), in our configuration the tilt contributes at most a minor, well-bounded systematic error and does not materially affect the fitted Young’s modulus.

**Figure S2.**
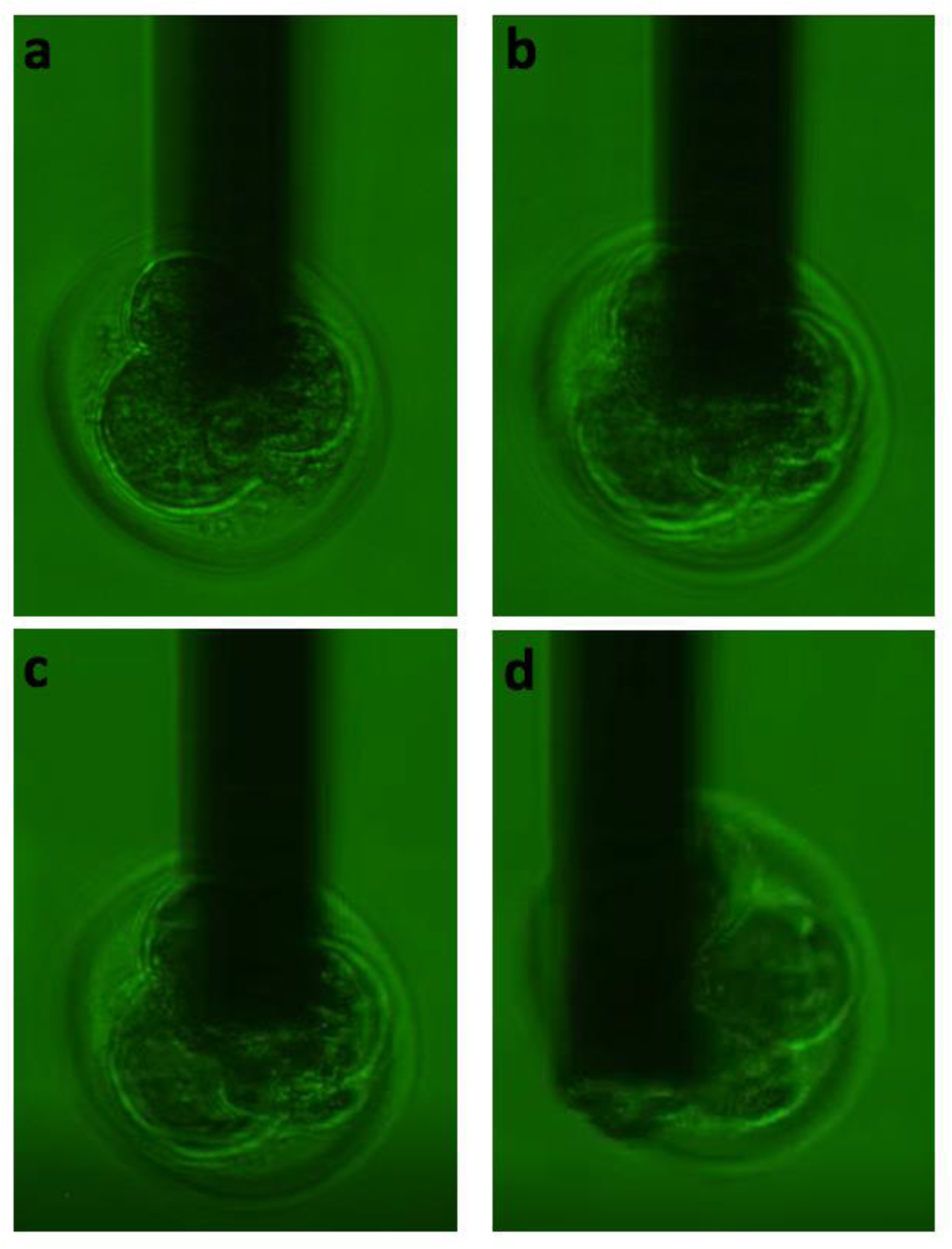
Time course during mechanical testing. **a)** At capture. **b)** After ∼5 minutes of measurements. **c)** ∼8 minutes after capture, a few seconds before release. **D)** Immediately after release. Bright-field images; same field of view.

**Figure S3.**
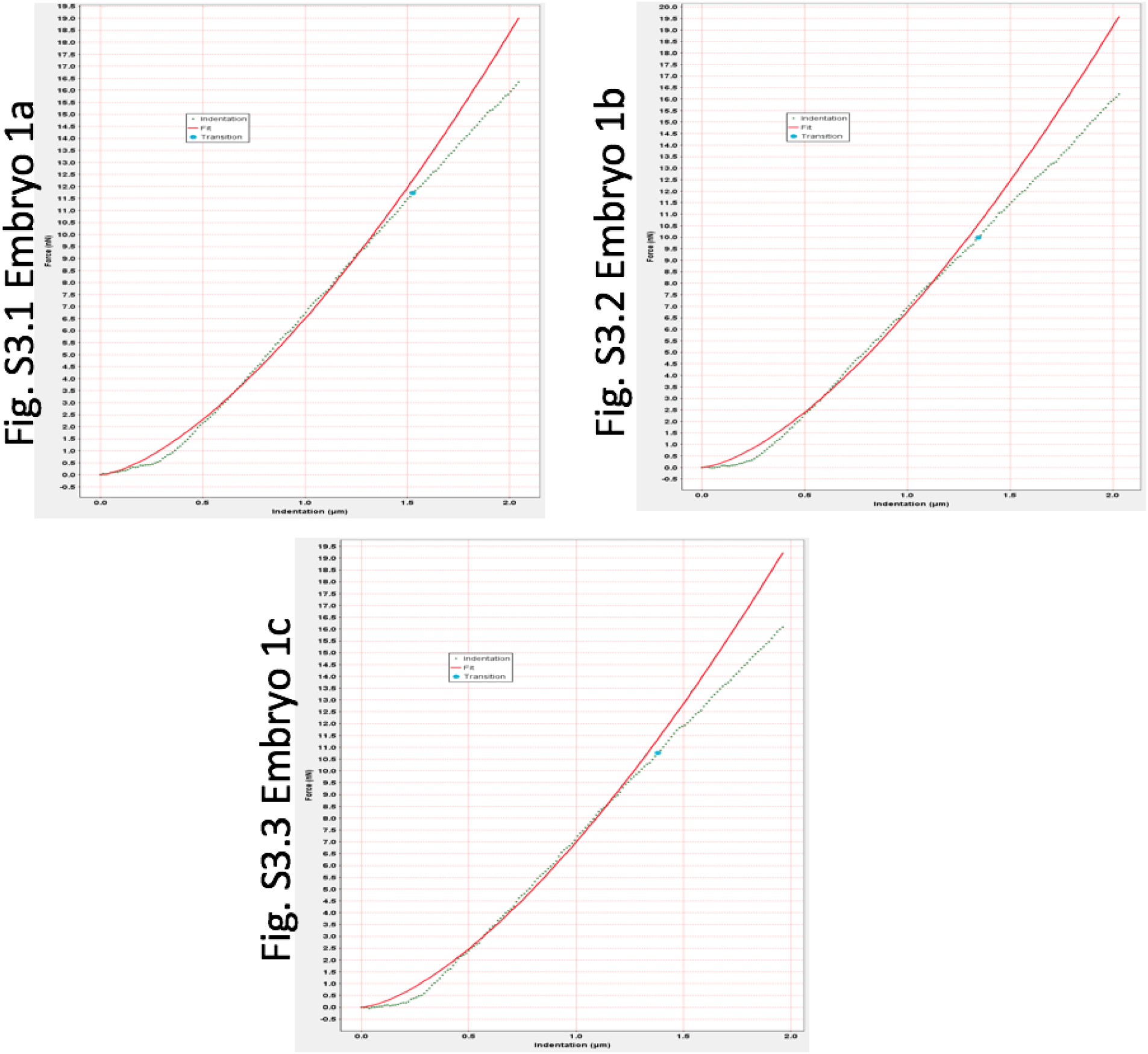
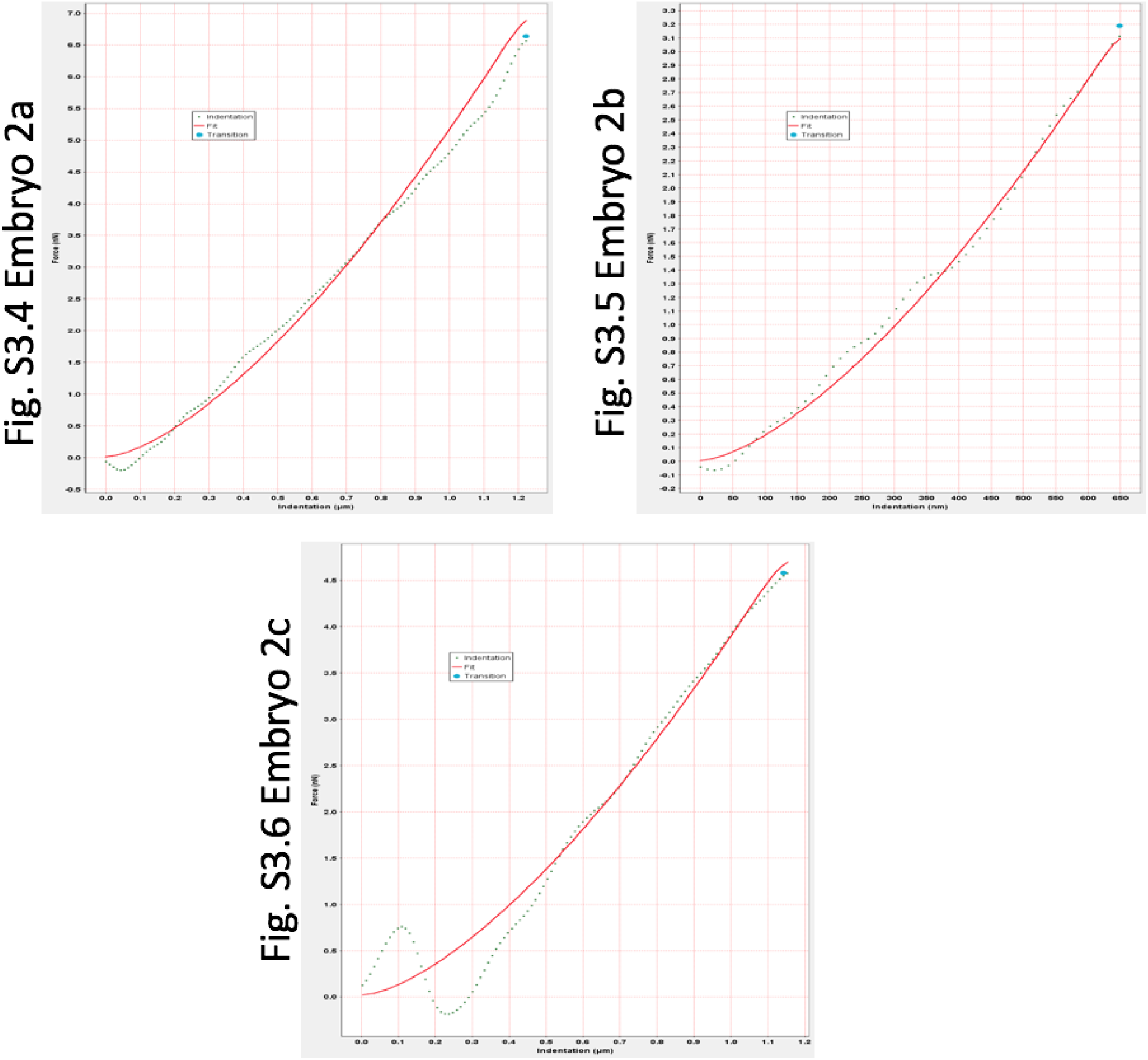
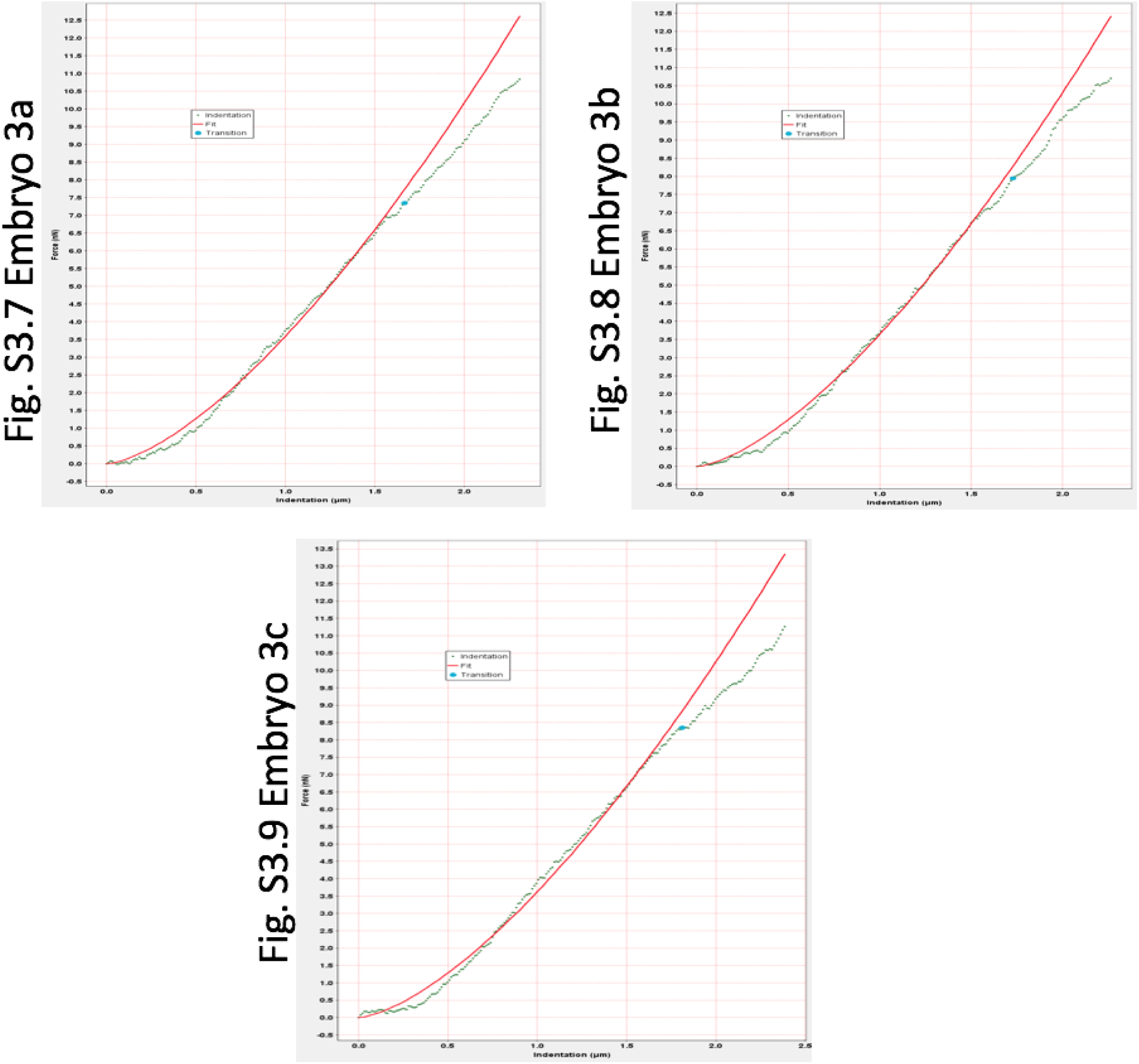
Indentation curves and Hertz fits. The dotted curve shows the raw indentation data; the red curve is the Hertz fit computed in AtomicJ (ν=0.5). The contact point (labeled on each panel) is identified automatically by AtomicJ’s model-based estimator. Plots were exported directly from AtomicJ.

**Table S1.** Young’s modulus and R^2^ values for each of the measurements. All analyses were performed in AtomicJ using the Paraboloid (Hertz) model (non-adhesive) with ν = 0.5. Across all nine curves the R² values are uniformly high (range 0.946–0.990, mean **≈** 0.968), indicating excellent agreement with the Hertz model in the selected fitting window.

| Measurement | Young's modulus (kPa) | R<sup>2</sup> |
| --- | --- | --- |
| Embryo 1a | 1.3661 | 0.9579 |
| Embryo 1b | 1.2696 | 0.9729 |
| Embryo 1c | 1.3226 | 0.9463 |
| Embryo 2a | 1.1718 | 0.9834 |
| Embryo 2b | 1.3614 | 0.9897 |
| Embryo 2c | 0.8823 | 0.9509 |
| Embryo 3a | 0.7752 | 0.9708 |
| Embryo 3b | 0.7863 | 0.98 |
| Embryo 3c | 0.782 | 0.9563 |

